# Females don’t always sing in response to male song, but when they do, they sing to males with higher pitched songs

**DOI:** 10.1101/860882

**Authors:** Alexander N. G. Kirschel, Zacharo Zanti, Zachary T. Harlow, Edgar E. Vallejo, Martin L. Cody, Charles E. Taylor

**Author notes:** Deceased 19^th^ June 2019 (Edgar E. Vallejo).

## Abstract

The long-held view that bird song is exclusively a male trait has been challenged recently by a number of studies and reviews highlighting the prevalence of female song. In spite of that, there remains a lack of knowledge on the function of female song, with most evidence thus far focusing on females performing duets with males in courtship displays, typically for joint territory defence or mate guarding purposes. Here we show in a tracheophone suboscine passerine *Formicarius moniliger*, a sexually monomorphic species in which both sexes sing, that females may participate in both intrasexual and intersexual territory defence. Females sing more in response to females than to males, suggesting they consider females more of a threat to their territory. Yet, females also demonstrate an unexpected pattern of singing back to playback of males singing higher frequency song than themselves. Unlike males, who respond indiscriminately to playback of any song performed by either sex, females appear to discern not only the sex, but perhaps also the size of the presumed intruder. There is a strong negative relationship between body mass and frequency, and females responding only to higher frequency male song suggests they will only engage in territory defence with males when they expect those males to be weaker than they are. While our results are consistent with expectations of a shared ancestral function of song in territory defence, they also suggest females may suffer greater costs in engaging in territorial disputes and thus limit their vocal contribution according to the perceived threat.

---

Sexual dimorphism is manifested in animals in a variety of ways resulting from different selective pressures on the sexes. Many elaborate traits have evolved, especially in males, with the aim of attracting females and fighting off rival males (Lande, 1980). Such dimorphism is consistent with conventional sex roles of males competing for mates and females investing in parental care (Kokko & Jennions, 2008). Yet many species are sexually monomorphic, with no differences in body size, colour or pattern. Sexual dimorphism may be constrained by genetic correlations or strong stabilizing selection acting on the sexes (Merilä et al., 1998) (Price & Grant, 1985), which could be driven by them occupying similar niches (Székely et al., 2007) or possibly biparental care (Badyaev, 1997). Many sexually monomorphic animals use acoustic signals for mate attraction and territory defence, and the traditional view has been that song is a sexually divergent behaviour performed by males, especially in temperate regions, but that view is now changing (Odom et al., 2014).

There is a growing number of studies on animals in which both sexes sing. It has long been known that in tropical birds, many females sing as well as males (Slater & Mann, 2004). Females in these instances may at times sing loosely in association with the male or at others sing a tightly-coordinated duet (Slater & Mann, 2004), the latter a phenomenon which has received much attention in the literature (Hall, 2004; Langmore, 1998; Slater & Mann, 2004). Female song in the tropics was previously considered a derived state attributed to sex role convergence in locations where birds are resident and defend territories year-round (Slater & Mann, 2004). However the view that it has resulted from a process of convergence in sex roles has been challenged by studies demonstrating female song is ancestral in songbirds (Odom et al., 2014; Riebel et al., 2019). Female song is indeed correlated with life history characteristics that favour female competition as found in many socially monogamous sedentary tropical birds (Price, 2009). It has been lost in clades that evolved differences in life history traits such as migratory behaviour and mating systems as the ranges of species in those clades shifted into temperate regions (Price et al., 2009). So while songs may represent ornaments attractive to the opposite sex, except in a few cases of sex-role reversal (e.g. Geberzahn et al., 2010; Goymann et al., 2004), their primary function in females is thought to be in intrasexual competition in species defending territories year-round from same-sex rivals (Tobias et al., 2011; Tobias et al., 2012b).

Such year-round territoriality is particularly evident in insectivorous birds (Hau et al., 2000; Slater & Mann, 2004; Tobias et al., 2011), with much previous work on duetting in birds focusing on insectivorous wrens and antbirds (Fedy & Stutchbury, 2005; Levin, 1985; Seddon & Tobias, 2006). Female song is also common in many Australian passerines, including in the insectivorous fairy-wrens, whistlers, shrike-thrushes, and bell birds, which like the wrens and antbirds, have extended longevity when compared to most temperate passerines, and defend territories year-round. In these taxa, the sexes are either sexually dimorphic or dichromatic (e.g. antbirds (Kirschel et al., 2019; Tobias & Seddon, 2009) and fairy-wrens and whistlers (Hall & Peters, 2008; van Dongen & Mulder, 2008)), sing distinctly different songs (e.g., wrens (Mennill & Vehrencamp, 2008)), or both (again antbirds (Tobias & Seddon, 2009), and fairy-wrens (Hall & Peters, 2008)), suggesting songs may still serve different functions among the sexes. Might there be situations in species where the sexes are monomorphic and sing indistinguishable songs, and what might such cases reveal regarding sex roles? Perhaps sexually monomorphic vocalisations could play a role in intersexual social competition (Tobias et al., 2012b). Here we examine such a case. We studied a population of sexually monomorphic Mexican antthrush – a.k.a. Mayan antthrush (Krabbe & Schulenberg, 2020) – (*Formicarius moniliger*), for which previous studies have shown both sexes sing (Kirschel et al., 2011; Kirschel et al., 2009b).

Suboscine passerines lack the vocal-control areas in the forebrain associated with song learning (Gahr, 2000) but exhibit some of the most elaborate mechanisms of sound production in their courtship displays, including wing stridulation in club-winged manakins (Bostwick & Prum, 2005)and aeroelastic flutter in *Smithornis* broadbills (Clark et al., 2016). Mexican antthrush is a tracheophone suboscine (Tobias et al., 2012a), however, a group whose simple stereotypic songs have allowed investigators to identify songs to species (Trifa et al., 2008), and to track movements of individuals in space and time using an acoustic sensor network (Collier et al., 2010). Previous work on Mexican antthrush has shown how songs can reliably be assigned to individuals (Kirschel et al., 2009a), and using such song classifications, resulting territory maps have demonstrated little territory overlap between same sex rivals suggesting strong intrasexual territoriality (Kirschel et al., 2011). However, previous studies have not distinguished male from female song (Kirschel et al., 2011) in a species which also lacks any sexual dimorphism in either plumage or body size (Krabbe & Schulenberg, 2020).

We nevertheless expected Mexican antthrushes could distinguish between the songs of each sex, and investigated how males and females respond to song produced by possible territory intruders of the same or opposite sex using playback experiments. We hypothesized that a vocal response to same sex playback represented territory defence, though could also represent mate guarding. Conversely, a vocal response to opposite sex playback could represent intersexual territory defence, but potentially also promiscuous intentions. Our overall aim was to determine when each sex used songs in response to possible intruders, thus informing us of the function of female song, which remains understudied across birds (Odom & Benedict, 2018; Riebel et al., 2019). Moreover, because of much evidence of a negative correlation between body mass and song frequency in birds (in centre frequency – (Wallschlager, 1980); and peak frequency (Ryan & Brenowitz, 1985), (but see (Geberzahn et al., 2009) we also tested whether such a relationship exists within antthrushes. If there is such a relationship, might individuals perceive differences in body mass? If larger birds sang lower frequency song, responses might differ based on the size of the presumed intruder. We tested this hypothesis by comparing responses to differences in frequency between responder’s song and playback stimulus, as well as differences in body mass between responder and presumed intruder.

## Methods

### Fieldwork

Fieldwork took place at the Estación Chajul in the Montes Azules Biosphere Reserve, in south-eastern Chiapas, Mexico (16°6′ 44″ N; 90°56′ 27″ W), during 8-17 June 2007, 7-18 December 2008, 7-28 May 2009, and 11-31 May 2012. Montes Azules, also known as Selva Lacandona, represents the largest expanse of pure tropical rainforest in North America (ParksWatch, 2003). The study was focused within a 50-ha study plot at an elevation range of 150-165 m, on the northern side of the Lacantún River (Kirschel et al., 2011). Mexican antthrush is a sedentary species (Krabbe & Schulenberg, 2020) that according to previous work defends a ~ 2ha territory year round, with both sexes participating in territory defence by singing, but do not coordinate their songs into duets (Kirschel et al., 2011). They are socially monogamous, and both sexes incubate eggs and provide parental care (corresponding author pers. obs.). Breeding has been recorded between April and June (Krabbe & Schulenberg, 2020), and our experience based on nest discoveries confirmed they were breeding in May upon the onset of the rainy season, but not in December.

We captured birds using target netting techniques and marked captured birds with a unique combination of colour rings to aid visual confirmation of their identity. We deployed individual mist nets (Avinet or Ecotone, 30- or 36-mm mesh, 12 × 2.5 m) along 1 - 2 m-wide trails, with the bottom of each net set at ground level. We target netted antthrushes using conspecific playback. Captured birds were also weighed and measured and 50-100μL of blood was obtained via venepuncture of the brachial vein for genetic sex identification. Individuals caught in 2012 also had a radio-frequency identification (RFID) tag fitted on their backs with a harness as part of a parallel study, and methods used are described therein (Corresponding author et al. unpublished ms).

We recorded ringed birds singing along trails in the study area in accordance with methods used in previous studies (Kirschel et al., 2011; Kirschel et al., 2009b). Specifically, we used a Marantz PMD 670 / 661 recorder at a sampling rate of 44 100/48 000 Hz and a 16-bit amplitude resolution, with Sennheiser microphones ME-67/K6, MKH20 microphone with a Telinga parabolic reflector, or MKH8050 housed in a Rycote windshield, as well as a VoxNet wireless acoustic sensor network recording at 48 000 16 bit samples per second (see (Collier et al., 2010; Kirschel et al., 2011).

### Ethical note

We minimised adverse impacts on the birds in procedures used, including keeping handling time to a minimum, ensuring bleeding had stopped after blood samples were taken, and returning birds for release back into their territories. In May 2012, we fitted 20 birds (11 males, 9 females) with RFID tags from BioTrack Ltd. (Dorset, UK; model PIP3) using a 1-mm-diameter elastic thread harness. We ensured the tag and harness were < 5% of body weight of all the birds involved, in accordance with recommendations (Fair et al., 2010). Antthrushes are sedentary, ground dwelling birds, and such loggers are likely less of a burden than for birds that migrate or forage above ground. The harness was designed to fall off after about a month, so that the bird carried the tag no longer than needed for the experiments. Fieldwork performed in June 2007, December 2008 and May 2009 was performed under UCLA’s Animal Research Committee protocol. Fieldwork in May 2012 was performed under a ringing licence from the Game and Fauna Service of the Republic of Cyprus. Fieldwork in Mexico were performed under SEMARNAT permit no. FAUT-0192.

### Playback experiments

Forty-one playback stimuli were prepared using Raven Pro 1.4 (Cornell Lab of Ornithology) using songs recorded in previous studies (Kirschel et al., 2011; Kirschel et al., 2009b) from 2007 to 2009, and further recordings obtained during fieldwork in 2012. The songs used belonged to 24 ringed individuals and had been recorded while colour-ring combinations were confirmed, or were classified unambiguously as belonging to specific individuals during previous work (Kirschel et al., 2011; Kirschel et al., 2009b). Sex of individuals whose songs were used in experiments had been previously identified genetically (Kirschel et al., 2011; Kirschel et al., 2009b), or had been predicted based on behaviour typical of one of the sexes during the May 2012 field season and subsequently confirmed genetically (see below). Each stimulus was prepared with songs that were high-pass filtered at 400 Hz to remove background noise, maximum amplitude normalized at 20 000 units (Raven’s amplitude unit), and arranged temporally with two songs of an individual within a stimulus with the remainder silence (approx. 22 s between songs) and then looped continuously for the duration of the experiment (see Table 1 for mean frequency and standard deviation values for male and female playback stimuli). Playback apparatus consisted of an Apple iPod MP3 player and a TivoliAudio PAL loudspeaker, with output volume set at the same level for all experiments. Playbacks were performed in December 2018, May 2009 and May 2012 in three sessions across the day and stimuli were randomly selected, except for the experiment subset in May 2012 (see below) and avoidance of neighbour song to minimise presentation of familiar stimuli (Lovell & Lein, 2004). Experiments were performed across the following times: during the morning session (0630 – 0841 hours), middle session (0914 – 1400 hours) and afternoon session (1523 – 1930 hours) within territory boundaries identified in the field (Kirschel et al. 2011).

**Table 1:** Mean (*X*) and standard deviation (SD) of peak frequency of male and female songs used in playback experiments and of the songs of individuals responding to them per field season.

| | Playback $\bar{X}$+SD | Response $\bar{X}$+SD |
| --- | --- | --- |
| Male song (Hz) | 2021.25+52.43 | 2018.16+43.62 |
| Female song (Hz) | 2021.7+64.66 | 2017.77+66.01 |

Experiments varied in length and depended on an approach by focal individuals. Antthrushes are inconspicuous, terrestrial birds (Cody, 2000), which may approach gradually and silently on the ground, possibly from far within their territory. In such cases it could take them 10 - 20 min, occasionally even longer, to approach to within a distance of the loudspeaker where a recording of suitable quality might be obtained. Recordings were thus obtained opportunistically when birds started vocalizing in response to the playback and only those experiments where recordings were obtained are included in analyses – except for a subset of experiments performed in May 2012 (see below). Our aim was to document who sang and what sex they were, in response to which individual playback and the sex, song characters, and body mass of the individual whose song was used in the experiment.

In experiments on pairs whose location was not known, if we had no response we dropped the experiment because we had no way of knowing if the pair were within range (c.f. (de Kort & ten Cate, 2001; Kirschel et al., 2009a). For some experiments, especially in field seasons prior to 2012, we may not have ringed both individuals of a focal pair. In such cases we still use the data of the experiment from the one identified individual but do not consider any partner’s response in analyses.

### May 2012 standardised experiments

A parallel telemetry study performed in May 2012 focused on approaches to specific stimuli (Corresponding author et al. unpublished ms). This set of experiments was arranged with a standardised playback duration of 20 min, and a balanced design of two male and two female playback experiments per territory. Playback experiments alternated in order per territory between Male-Female-Male-Female and Female-Male-Female-Male with one experiment in the morning session (from 0810 – 0830 hours), two in the middle session (from 1110 - 1130 hours and 1340 – 1400 hours) and one in the afternoon session (from 1740 – 1800 hours). We waited at least 24 hours before performing the next experiment in any territory. We set the duration to 20 min because using telemetry we could determine the locations of both members of the pair before the initiation of playback. Because these experiments had a cut-off time, we also kept track of experiments where neither individual of a pair responded with song within that time and included them in our analyses.

### Genetic sex identification

DNA was extracted from blood or feather samples collected from 21 individuals in the field in 2012 using a QIAGEN DNEasy Blood and Tissue Kit (QIAGEN, Valencia, CA) following manufacturer’s protocols. Highly conserved primers (2550F and 2718R, (Fridolfsson & Ellegren, 1999)) were used in polymerase chain reaction (PCR) to amplify the differently sized introns of Z- and W-linked chromohelicase-DNA binding protein 1 (CHD1) genes. PCR products were separated in a 1 or 2% agorose gel run in TAE buffer, revealing one or two bands for males and females respectively.

### Song feature extraction

Song features needed to be extracted from recordings for use in analyses. These included the peak frequency measurements needed to determine if song frequency affects response levels. Also, to identify the singer on recordings obtained in 2012, we needed to extract song features from recordings and then classify songs to individual in a canonical discriminant analysis, as had been done with recordings from previous years (Kirschel et al., 2011). We thus imported 83 recordings from fieldwork performed in May 2012 into RAVEN Pro 1.4 (Cornell Lab of Ornithology, Ithaca, NY, USA) where song sequences were cut into separate WAV files, which were then processed in a feature extraction program in MATLAB 7 (MathWorks 2005). The procedure has been described in detail previously (Kirschel et al., 2009b). Here, we follow the approach used in a previous study (Kirschel et al., 2011), extracting 19 temporal and spectral features of song: Temporal features extracted were (1) inter-onset-interval of the first note to the second note; (2) duration from the start of the second note to the end of the last note; (3) rate of the main trill part of the song; (4) rate of the first half of the trill; (5) rate of the second half of the trill (see Kirschel et al. 2009a) for an explanation of rate calculations used), and (6) number of notes. We extracted the following spectral measures: the peak frequency of the first five, last four, and the middle note of the trill; the highest and lowest frequency notes; and the peak frequency of the whole trill.

### Differentiation of multiple individuals singing on a single recording

Where two or more singing individuals were recorded on any recording, we separated out the songs to each individual first based on any instructions provided by the recordist (e.g. announcements of bird east of trail, west of trail following each song), and/or any obvious differences between songs observed from spectrograms which allowed for clear separation of which songs were sung by each bird. In such a case, the threshold of 70% assignment of songs from a recording to a single individual was applied separately to each individual singing on each recording. Likewise, the playback stimulus was typically also on recordings and observers recognised the playback on the recording and ensured they did not extract features from it.

### Canonical discriminant analysis

In order to identify the singers responding to playback experiments and to extract song frequency values for analyses, we needed to identify them even when we could not see their colour ring combinations (if ringed). To do this we classified songs from recordings to individuals. The identity of the singer in 168 cases out of 230 identified on recordings had been determined in previous studies (Kirschel et al., 2011; Kirschel et al., 2009b) or by sight or with radio telemetry in 2012. However, there remained 39 recordings of antthrush songs for which we still needed to identify the singer. There are numerous ways to classify songs to individuals, which can include supervised and unsupervised methods (Blumstein et al., 2011), and several such methods have been tested on songs of Mexican antthrush (Kirschel et al., 2009b).

We followed (Kirschel et al., 2011) in using linear canonical discriminant analysis (CDA) to identify songs to individuals. The CDA was performed in STATA 11.2 (StataCorp 2009) on songs cut from recordings obtained in May 2012 using the 19 variables extracted from the songs. Following (Kirschel et al., 2011), we used 1 189 songs from recordings where the singer could be unambiguously confirmed by sight or radio telemetry as training data and predicted the singer of 619 songs on the remaining 39 recordings.

We then calculated the percentage correct classification based on the number of songs from a recording classified to the same individual and compared the classification with expectations based on the location of the recording. If the individual identified as the singer was one of the focal pair or an individual from a nearby territory, we assumed that the individual on the recording was correctly identified if at least 70% of the songs were assigned to that individual, in accordance with Kirschel et al. (2011). Following the CDA classification, we filtered out the recordings with less than 70% assignment to one identified individual except for any recordings where either the individual was identified in the field either by sight or using radio telemetry (corresponding author et al. unpubl. ms).

The resultant songs classified to individual were then pooled with the 2011 songs from 168 individuals on recordings previously classified (Kirschel et al., 2011; Kirschel et al., 2009b), and statistical analyses were performed on a total of 3022 songs attributed to 24 individuals. Moreover, we tested whether with the combined dataset of 50 individuals we could distinguish between male and female song in a CDA with leave-one-out-classification using the mean values of the 19 song features from the entire set of recordings included in the study.

### Statistical analyses

To determine whether there was a relationship between body mass and song frequency we first calculated the mean peak frequency from all recordings of each individual for each field season. Peak (or dominant) frequency has been shown to be strongly negatively correlation with body size in the closely related suboscine passerine group, the woodcreepers (Torres et al., 2017). For this analysis, we calculated the mean peak frequency only from recordings where the bird’s ring combination was identified, or where the bird was identified using radio telemetry (Corresponding author et al. unpublished ms). Body mass varies between seasons, especially in females. We controlled for seasonal variation by using values per season for each individual in analyses (and mean values per season if recaptured). We then ran a GLMM in lme4 in R, fitted with a Gaussian distribution, with mean peak frequency as the dependent variable and body mass (g) and sex as fixed effects and individual included as a random effect, to account for individuals recorded (and body mass measured) over several years.

We then tested whether one of the sexes sings more than the other in response to playback of either sex using a chi-squared test. We restricted this analysis to 167 experiments (83 male and 84 female song playbacks) to 32 territorial individuals in pairs where both members of the pair were ringed. To test for what determines when each sex sings, we used Generalized Linear Mixed Models (GLMM) built in ‘glmmTMB’ in R, fitted with the binomial distribution and logit link function. We determined the effect on the binary response variable “female song response” or “male song response” of the difference in peak frequency between the mean peak frequency of the responder’s song (from all recordings of the individual obtained that year) and the peak frequency of the playback stimulus (hereafter frequency differential). We log-transformed all frequencies because doing so best approaches the scale with which animals perceive and modulate their frequencies (Cardoso, 2013). We also tested for an effect of the difference in body mass between subject and playback singer (hereafter body mass differential) and the latter’s sex affected male or female response levels. Body mass was also log-transformed, because log-size is linearly related to log frequency (Torres et al., 2017). We included in the model the binary predictor of sex of the individual whose song was used in the playback experiment, and its interaction with frequency and body mass differentials.

Responses to song playback might also be affected by the season, the time of day, and whether the bird had previously been captured in a mist net in response to playback. We thus included season, capture history and session as fixed factors in the model. Capture history was defined as a categorical factor with the following categories: 1) never caught before; 2) last caught in a previous field season (i.e. over five months earlier); 3) caught in current field season. Because responses could also vary by individual subject’s propensity to sing in response to playback, or because of other features of a specific stimulus used, we included subject of experiment and playback stimulus used as crossed random factors in the models. While we controlled for session in our models, we also tested whether there was any bias towards playing more playbacks of a certain sex at a time when subjects were more or less likely to respond by performing a chi-squared test.

Using the subset of 36 experiments performed with a standardised methodology in May 2012, we tested using GLMMs in ‘lme4’ in R specific questions on how frequency and body mass differentials may explain when males and females responded to playback. Specifically, we tested whether frequency (or body mass) differential varied between the male/female playback experiments that subjects (of each sex) did and did not respond to (with territory as a random factor).

We used the DHARMa package in R to check the residual diagnostics of our models, inspecting the QQ plot and confirming there was no significant deviation evident from the Kolmogorov-Smirnov test.

## Results

### Genetic sex identification

Of the 21 birds we obtained samples from in 2012, 13 were male and 8 were female. This information was combined with the data from previous genetic analyses (Kirschel et al., 2011), from which 12 sexed males and 11 females, were included as playback stimuli singers (13 male, 11 female) and/or test subjects (22 male, 17 female) in playback experiments (i.e. the 39 birds included in experiments whether they responded with song or not).

### Canonical discriminant analysis

Based on the CDA trained using songs of 20 individuals, 619 songs from 39 recordings were classified. Of these, 88.7% were assigned to the predicted individual, consistent with classification rates found previously for similar numbers of individuals in a season (Kirschel et al., 2011). By contrast, the CDA could only classify 75.1% of male song and 55.9% of female song to the correct sex (See Table S2 for CDA coefficients and canonical structure).

### Patterns of variation in body size and song frequency

While there was no difference in (log) peak frequency according to sex (Table 2, see also Table 1) there was a significant negative relationship between body mass and peak frequency, meaning larger individuals sang lower frequency songs (Fig. 1), though there was no effect of season, nor an interaction effect of sex and body mass on song frequency.

**Figure 1.**
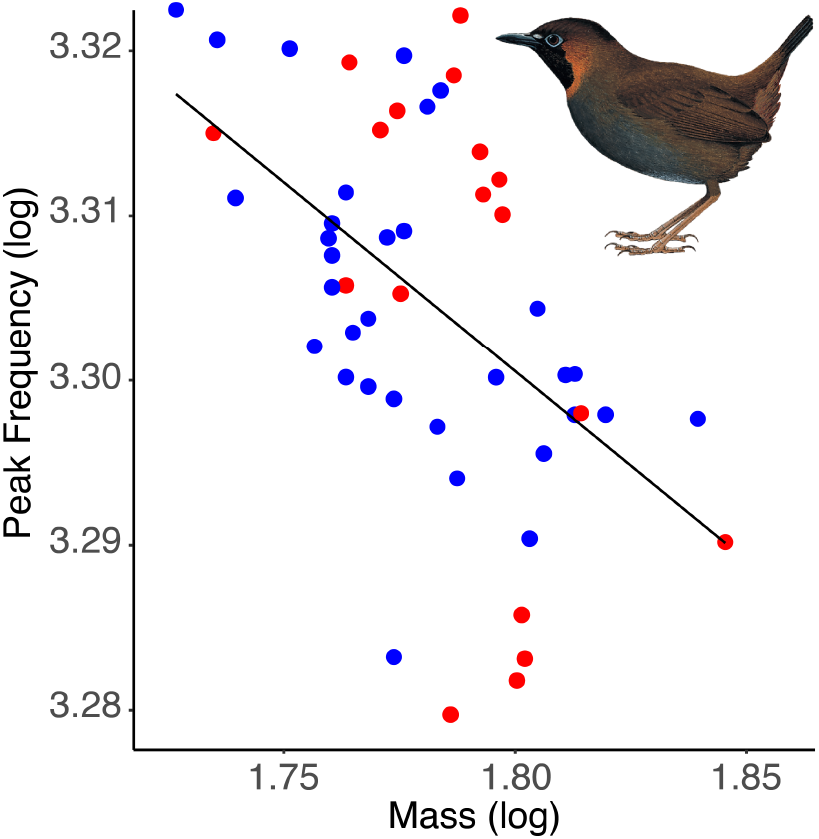
Relationship between (log) body mass and (log) peak frequency in males (blue dots) and females (red dots). *Formicarius moniliger* illustration courtesy of (Krabbe & Schulenberg, 2020).

**Table 2:** GLMM with Gaussian distribution testing for the effect of mass, sex, and their interaction, as well as season, on peak frequency (log), with individual as a random factor.

| N = 50, individuals = 40 | Estimate | SE | t | P |
| --- | --- | --- | --- | --- |
| Intercept | 3.75 | 0.14 | 27.70 | <0.0001 |
| Sex | 0.07 | 0.23 | 0.31 | 0.76 |
| Body mass (log) | -0.25 | 0.08 | -3.26 | 0.002 |
| Season | -0.001 | 0.002 | -0.63 | 0.54 |
| Sex/body mass interaction | -0.04 | 0.14 | -0.29 | 0.77 |

### Playback experiments

Males responded significantly differently to females, responding more with song to both male (88.2% vs 11.8% of experiments; *χ*^2^ = 5.34, *P* = 0.02) and female playback (88.0% vs 27.7% of experiments; *χ*^2^ = 15.52, *P* < 0.0001). Males showed no difference in their likelihood to sing based on sex of playback, frequency or body mass differentials or their interactions with sex (Table 3, Fig. 2). By contrast, females were more likely to respond with song to female playback than male playback, but not in relation to frequency or body mass differentials (Table 3). There was, however, a significant negative interaction between sex of playback and frequency differential, indicating that females increasingly respond to male song the lower their own song was in frequency compared to the male playback stimulus (Fig. 2). There was no interaction effect of sex with body mass differential though, and no effect of season or capture history on responses either (Table 3), but females were less likely to respond in the middle and afternoon sessions. There was, however, no bias regarding which session male and female playbacks were presented in (*χ*^2^ = 0.48, *P* = 0.79).

**Figure 2.**
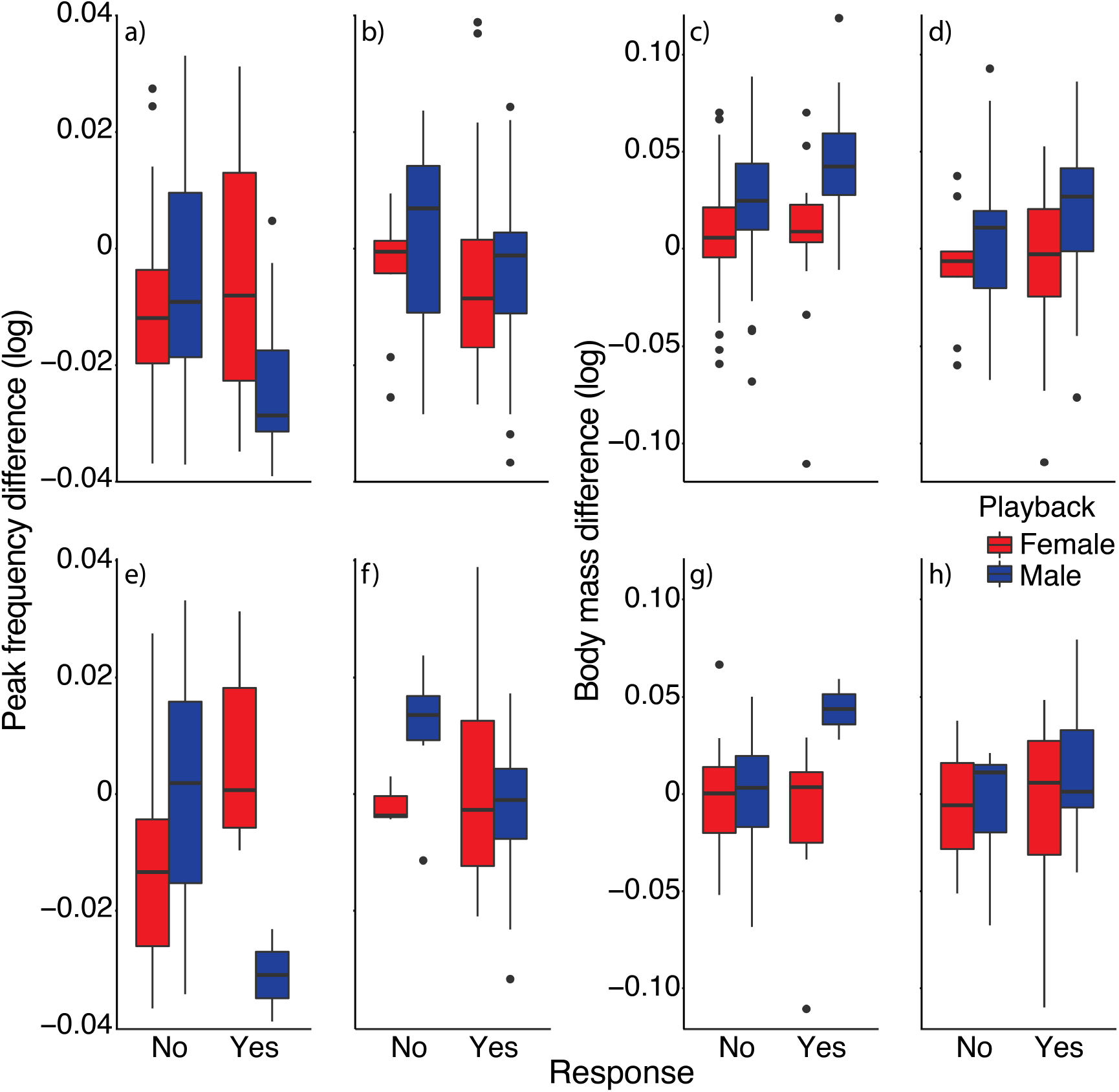
Female (a) and male (b) peak frequency difference with male / female playback stimulus (frequency differential) according to sex of playback stimulus singer in experiments they do and do not respond to (with song). Female (c) and male (d) body mass difference with male / female playback stimulus singer (body mass differential) in experiments they do and do not respond to. Female (e) and male (f) frequency differentials according to sex of playback stimulus singer in experiments they do and do not respond to from subset of experiments performed in May 2012. Female (g) and male (h) body mass differentials in experiments they do and do not respond to from subset of experiments performed in May 2012. Lower (more negative) frequency differentials represent individuals singing in response to higher frequency song then their own. Higher (positive) body mass differentials represent individuals responding to playback stimulus singers that have lower body mass then they do.

**Table 3:**
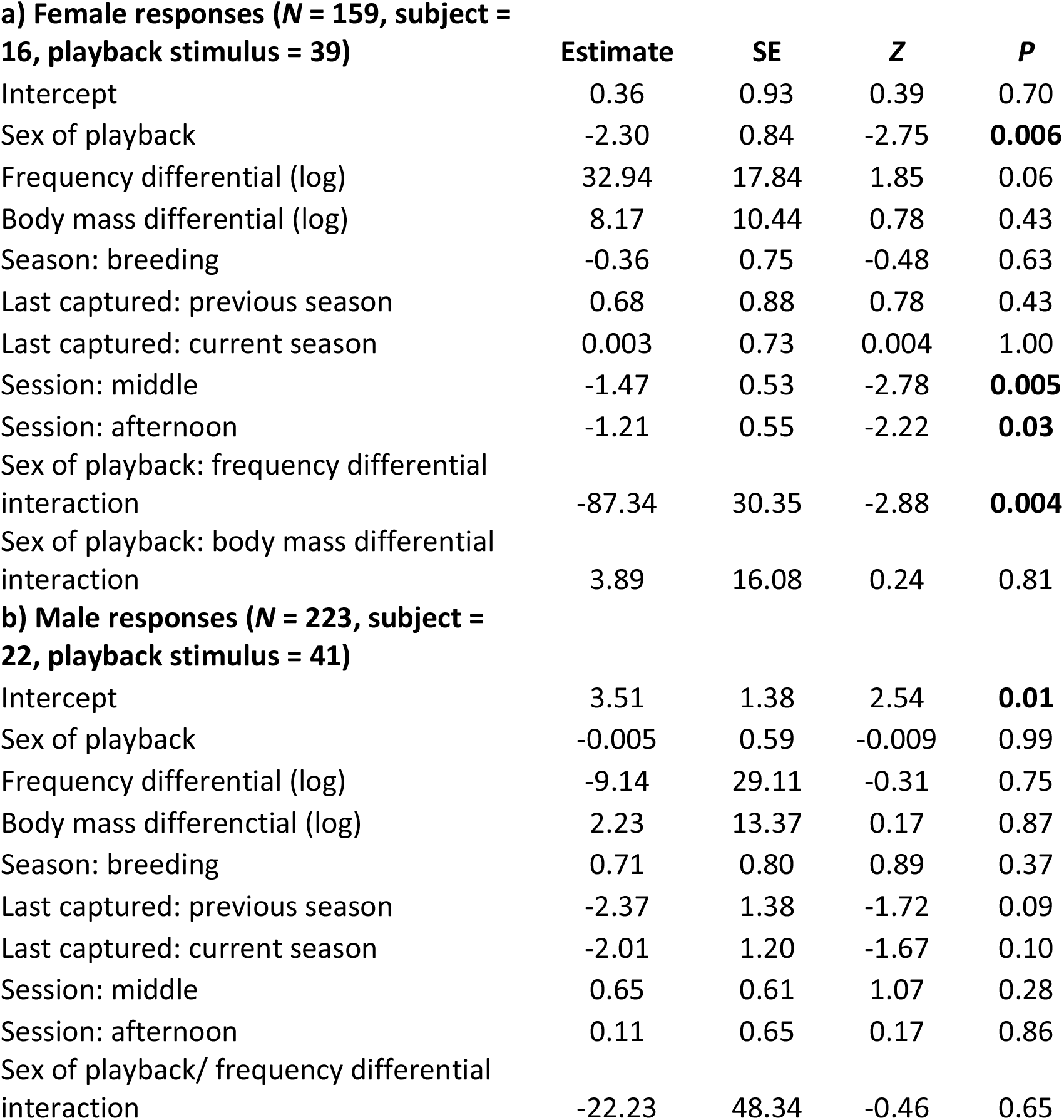
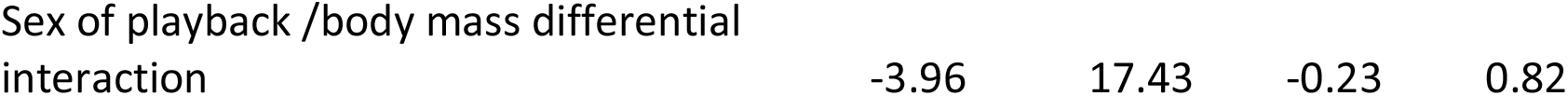
Results of GLMMs testing the effect of sex of playback, frequency and body mass differential, and the interaction of sex of playback with those differentials on vocal responses of (a) females and (b) males.

|  | **Estimate** | **SE** | **Z** | **P** |
| --- | --- | --- | --- | --- |
| Intercept | 0.36 | 0.93 | 0.39 | 0.70 |
| Sex of playback | -2.30 | 0.84 | -2.75 | **0.006** |
| Frequency differential (log) | 32.94 | 17.84 | 1.85 | 0.06 |
| Body mass differential (log) | 8.17 | 10.44 | 0.78 | 0.43 |
| Season: breeding | -0.36 | 0.75 | -0.48 | 0.63 |
| Last captured: previous season | 0.68 | 0.88 | 0.78 | 0.43 |
| Last captured: current season | 0.003 | 0.73 | 0.004 | 1.00 |
| Session: middle | -1.47 | 0.53 | -2.78 | **0.005** |
| Session: afternoon | -1.21 | 0.55 | -2.22 | **0.03** |
| Sex of playback: frequency differential interaction | -87.34 | 30.35 | -2.88 | **0.004** |
| Sex of playback: body mass differential interaction | 3.89 | 16.08 | 0.24 | 0.81 |
| **b) Male responses (*N* = 223, subject = 22, playback stimulus = 41)** |  |  |  |  |
| Intercept | 3.51 | 1.38 | 2.54 | **0.01** |
| Sex of playback | -0.005 | 0.59 | -0.009 | 0.99 |
| Frequency differential (log) | -9.14 | 29.11 | -0.31 | 0.75 |
| Body mass differential (log) | 2.23 | 13.37 | 0.17 | 0.87 |
| Season: breeding | 0.71 | 0.80 | 0.89 | 0.37 |
| Last captured: previous season | -2.37 | 1.38 | -1.72 | 0.09 |
| Last captured: current season | -2.01 | 1.20 | -1.67 | 0.10 |
| Session: middle | 0.65 | 0.61 | 1.07 | 0.28 |
| Session: afternoon | 0.11 | 0.65 | 0.17 | 0.86 |
| Sex of playback/ frequency differential interaction | -22.23 | 48.34 | -0.46 | 0.65 |
| Sex of playback /body mass differential<br>interaction | -3.96 | 17.43 | -0.23 | 0.82 |

In the subset of experiments performed systematically in May 2012, we found that male songs females responded to were significantly lower in frequency compared to their own song than the male songs they did not respond to were (Fig. 2), but there was no significant difference found in the other tests of frequency differentials (i.e., females or males responding to female song, or males responding to male song) (Table S3). Further, we found that the males whose songs females responded to were much lighter in comparison to their own mass than the males they did not respond to were (Fig. 2). Again, there was no significant pattern in the remaining tests relating to body mass differentials within and among the sexes that subjects did and did not respond to (Table S3).

## Discussion

In this study, antthrush females responded with song to playback less often than males did, supporting the view that males devote more effort to territory defence than females do. Indeed, males appeared to respond indiscriminately to song, whether performed by males or females and irrespective of the body size of the presumed intruder. Such responses suggest either heightened aggression towards intruders of both sex in males because they pose a similar threat, promiscuous intentions when females trespass into their territories, or an inability to distinguish song to sex. By contrast, females responded significantly more often to female song than to male song, suggesting they do discriminate between the songs of each sex, in spite of the extent of overlap in song characteristics (Fig. 1), and our failure to reliably classify songs to sex in a discriminant analysis. Thus, signals that may appear sexually monomorphic to human observers are still likely to contain information on sex-specific differences (Price, 2015). Females singing in response to female song is consistent with both the function of intrasexual territory defence and mate guarding (Cain & Langmore, 2015; Langmore, 1998; Levin, 1996; Tobias & Seddon, 2009). Mexican antthrushes are territorial year-round and form long-term pair bonds and we believe our results here, supported by evidence from radio telemetry and field observations (corresponding author unpublished ms), suggest females guard their mates from potential rival females. Yet, on occasion, females responded to male song. They responded significantly more to male song the lower the frequency their song was compared to the male song. In other words, they respond to those males they perceived as singing higher pitched songs than they did.

Peak frequency has been shown to be negatively correlated with body size in birds, both among species (Ryan & Brenowitz, 1985), and within species (Hall et al., 2013). Here, we tested whether there was a relationship between peak frequency and body mass in individuals of Mexican antthrush. We found that larger, heavier antthrushes do indeed sing lower frequency songs than lighter antthrushes, while song frequency does not differ between the sexes. What this relationship suggests is that females responding to males singing at higher frequencies are responding to smaller males. We did not find an overall effect of body size differential on female responses, though females did sing in response to smaller males than those they did not respond to in the systematically designed subset of experiments in May 2012. Of course, there is variation in song frequency in animals that is not explained entirely by body mass (Fitch, 1999), and the antthrushes responded to what they could hear and not what they could see. Other aspects of the playback stimulus might provide further information to the receiver regarding the size of the singer and the threat (or opportunity) represented. One possibility is that individuals may be able to lower their frequency as a signal of aggressive intent (e.g. Geberzahn et al., 2010). However, the consistency of spectral and temporal features that results in the levels of classification accuracy achieved for the species, even between years (Kirschel et al., 2011), suggests song frequency is unlikely to vary according to social context.

We acknowledge that our experiments were not standardised for duration during earlier field seasons when we also did not keep track of experiments with no response at all, although our standardised experiments performed in May 2012 provided consistent results. The 2012 experimental subset demonstrated both a greater frequency, and a greater body mass, differential between subject and singer in the male playback experiments females responded to than those they did not respond to, providing further indirect as well as direct support that females responded to smaller males. Might females be attracted to males that are smaller than them? Or could responses to smaller males just be an indirect consequence of a preference for higher frequency song (see Cardoso, 2012)? We suggest an alternative explanation. These paired females are not responding to male song for mutual attraction purposes. Instead, we believe that females will participate in intersexual territory defence by singing back only when they perceive the intruding male to be smaller, and thus weaker, than they are. Sexually monomorphic songs are expected to fulfil a similar function between the sexes (Riebel et al., 2019), and thus could be used for both intrasexual and intersexual territory defence. But in a species where the female sings a fraction of what the male does, it seems she chooses carefully when she will use her song for the purpose of intersexual territory defence. Indeed, females did not discriminate between the frequencies of female songs, which coupled with their significantly higher likelihood to respond to female than male song suggests they see females as more of a threat to their territory than males. But the cost of intersexual territory defence could be especially high against larger males (Logue & Gammon, 2004), so they respond only to those males they believe they have a physical advantage over.

We did not test whether female song was influenced by the male mate’s response and thus formed a coordinated territorial response to intruders (e.g., (Hall & Peters, 2008)– males responded to almost every experiment rendering such a test unworkable. Such coordinated territorial responses are typically found in birds that arrange their songs into duets, which antthrushes do not do. Although we have not tested for it here, our observations suggest that female antthrushes do not sing in tandem with the male or jam his song as a mate guarding strategy, as found in duetting *Hypocnemis* antbirds (Tobias & Seddon, 2009). Instead, females sometimes sing solo (see also Kirschel et al., 2011). We also caution that our results are based on a large number of recordings of individuals identified based on song classifiers, and that we excluded songs on recordings that we were unable to classify to specific ringed individuals. Such songs might even have belonged to female or male partners of the individuals that we did identify singing on recordings, but they were not included in analyses if the CDA failed to identify them. Nevertheless, any situations where we were unable to identify the individual singer reduced our sample size and overall statistical power. We have no reason to assume any missing data would not be representative of the patterns reported here. We also caution that nonvocal responses may play an important part in both territorial defence and mate guarding, so it would be important to associate nonvocal responses such as approaches to playback stimuli as well as vocal responses.

Many previous studies on female song in Neotropical suboscines have shown it to be dimorphic from the male song and are typically coordinated into duets (e.g. Bard et al., 2002; Fedy & Stutchbury, 2005; Roper, 2005; Seddon & Tobias, 2006) consistent with divergent functions between the sexes. We have shown here that female antthrush songs cannot reliably be distinguished from male song by human observers and classifiers. We do not suggest, however that the similarity of female song with male song is evidence for convergence in sex roles. Instead, we suggest our study on suboscine passerines is consistent with the premise that female song is ancestral in songbirds (Odom et al., 2014; Riebel et al., 2019) with females singing to defend territories much in the way that males do. Nevertheless, we find that females respond to song playback far less than males do and pick and choose when they will respond with song to presumed intruders, especially when they are males.

