## Supplementary material for "Females don’t always sing in response to male song, but when they do, they sing to males with higher pitched songs"

**Contents**

Data table S1

Statistical tables S2, S3

**Table S1:** Individuals ringed at Estacion Chajul incorporating those included in experiments as playback stimulus singers and/or playback subjects identified by Band (Colour-ring combination), date caught in each field season, sex, mass when caught and mean peak frequency of their songs that year from recordings where they were seen or identified through radio telemetry. The list includes some individuals that were ringed but not included in experiments.

| Band | Date caught | Year | Sex | Mass(g) | Peak frequency | Singer/ subject/ both |
|---|---|---|---|---|---|---|
| BOY | 18-May-12 | 2012 | Male | 59.20 | 2035.63 | both |
| BRO | 16-May-12 | 2012 | Male | 60.70 | 1982.31 | both |
| BUB | 14-May-12 | 2012 | Female | 54.30 | 2065.43 | both |
| BUG | 19-May-12 | 2012 | Male | 58.00 | 1996.09 | subject |
| BYK | 19-May-12 | 2012 | Male | 59.40 | 1919.68 | both |
| BYR | 20-May-09 | 2009 | Female | 62.60 | 2052.11 | subject |

| | | | | | | |
|---|---|---|---|---|---|---|
| CPG | 09-Dec-08 | 2008 | Female | 63.00 | | |
| CPG | 11-May-09 | 2009 | Female | 70.05 | 1950.65 | both |
| CWC | 09-Dec-08 | 2008 | Male | 54.50 | | |
| CWC | 07-May-09 | 2009 | Male | 53.30 | 2101.28 | both |
| GGG | 29-May-12 | 2012 | Male | 58.00 | 2048.43 | neither |
| GGR | 10-Jun-07 | 2007 | Male | 65.00 | 1985.64 | |
| GGR | 14-Dec-08 | 2008 | Male | 69.10 | 1984.49 | |
| GGR | 12-May-09 | 2009 | Male | 66.00 | 1985.65 | |
| GGR | 11-May-12 | 2012 | Male | 64.70 | 1996.68 | both |
| GOB | 12-May-12 | 2012 | Male | 54.40 | 2092.50 | both |
| GRU | 11-Dec-08 | 2008 | Male | 58.65 | 2012.68 | |
| GRU | 16-May-09 | 2009 | Male | 58.65 | 1993.47 | both |
| GUY | 13-May-12 | 2012 | Female | 62.10 | 2047.85 | subject |
| GWW | 08-Jun-07 | 2007 | Male | 57.50 | 2035.37 | |
| GWW | | 2008 | Male | 57.60 | 2039.60 | |
| GWW | 09-May-09 | 2009 | Male | 57.60 | 2030.52 | both |
| GYR | 25-May-12 | 2012 | Female | 63.30 | 1930.96 | subject |
| KOW | 12-May-12 | 2012 | Male | 63.80 | 2015.46 | subject |
| KRG | 09-Jun-07 | 2007 | Female | 61.20 | 2082.17 | singer |
| KUG | 09-May-09 | 2009 | Female | 59.30 | | neither |
| KYC | 10-Jun-07 | 2007 | Male | 61.30 | 1968.09 | |
| KYC | 12-Dec-08 | 2008 | Male | 62.30 | | |
| KYC | 10-May-09 | 2009 | Male | 62.50 | 1996.05 | both |
| ORK | 16-May-12 | 2012 | Male | 60.00 | | neither |
| OUO | 11-May-12 | 2012 | Male | 60.40 | 2073.05 | both |

| | | | | | | |
|---|---|---|---|---|---|---|
| PUB | 20-May-09 | 2009 | Male | 59.70 | 2087.90 | both |
| PUU | 07-Dec-08 | 2008 | Male | 57.10 | 2004.79 | |
| PUU | 13-May-09 | 2009 | Male | 57.60 | 2021.49 | both |
| PWG | 13-Dec-08 | 2008 | Female | 60.40 | | neither |
| PYC | 17-May-09 | 2009 | Male | 65.00 | 1996.97 | both |
| RGR | 21-May-09 | 2009 | Male | 60.80 | 2077.78 | both |
| RKW | 10-Jun-07 | 2007 | Female | 72.40 | | |
| RKW | 14-Dec-08 | 2008 | Female | 62.00 | 2060.09 | both |
| ROO | 13-May-12 | 2012 | Female | 61.40 | 2099.61 | subject |
| ROY | 12-May-12 | 2012 | Male | 59.20 | | neither |
| RPC | 10-Dec-08 | 2008 | Female | 57.60 | | |
| RPC | 14-May-09 | 2009 | Female | 59.50 | 2071.90 | both |
| RRP | 08-May-09 | 2009 | Female | 59.60 | 2019.69 | both |
| RRR | 04-Feb-10 | 2010 | | 63.30 | | neither |
| RRR | 24-May-12 | 2012 | Male | 56.40 | 2089.94 | both |
| RWB | 21-Jan-10 | 2010 | | 57.50 | | neither |
| RYY | 09-Jun-07 | 2007 | Female | 67.00 | | |
| RYY | 26-May-09 | 2009 | Female | 62.70 | 2042.12 | both |
| UCY | 08-Dec-08 | 2008 | Female | 54.60 | | |
| UGW | 19-May-09 | 2009 | Male | 59.40 | 1990.03 | both |
| UUW | 15-May-09 | 2009 | Female | 52.40 | | |
| WKW | 11-Dec-08 | 2008 | Female | 63.40 | 1919.21 | |
| WKW | 17-May-09 | 2009 | Female | 63.15 | 1913.38 | both |
| WKW | 19-May-12 | 2012 | Female | 62.40 | | |
| WOW | 15-May-12 | 2012 | Female | 61.10 | 1904.30 | both |

| | | | | | | |
|---|---|---|---|---|---|---|
| WPB | 12-May-09 | 2009 | Female | 59.40 | | |
| WPB | 12-May-12 | 2012 | Female | 61.80 | | subject |
| WRR | 15-Jun-07 | 2007 | Female | 55.50 | | neither |
| WRY | 15-May-12 | 2012 | Male | 64.00 | 1974.89 | both |
| WYG | 19-May-12 | 2012 | Female | 58.00 | 2022.04 | both |
| YBY | 12-May-09 | 2009 | Female | 62.60 | | neither |
| YKK | 23-May-09 | 2009 | Male | 54.90 | 2046.85 | subject |
| YOU | 29-May-12 | 2012 | Male | 60.00 | | |
| YRU | 21-Jan-10 | 2010 | Female | 56.00 | | |
| YRU | 13-May-12 | 2012 | Female | 58.10 | 2085.94 | both |
| YUK | 16-May-12 | 2012 | Male | 58.20 | 2008.74 | |
| YUW | 09-Dec-08 | 2008 | Female | 64.00 | | |
| YUW | 09-May-09 | 2009 | Female | 65.20 | 1986.05 | subject |
| YWR | 10-Jun-07 | 2007 | Female | 58.70 | | |
| YWR | 16-May-09 | 2009 | Female | 59.00 | 2066.34 | subject |
| YYG | 16-Jun-07 | 2007 | Male | 59.70 | 2037.43 | |
| YYG | 10-Dec-08 | 2008 | Male | 68.00 | | |
| YYG | 14-May-09 | 2009 | Male | 63.55 | 1951.62 | both |

**Table S2**: Standardised canonical discriminant function coefficients and canonical structure of function 1.

| Song Feature | coefficients | structure |
|---|---|---|
| trill rate | -1.72 | -0.16 |
| rate first half | -0.79 | -0.17 |
| rate second half | -0.25 | -0.13 |
| No. of notes | 5.3 | -0.03 |
| IOI note 1 to note 2 | 0.33 | 0.35 |
| Trill duration | -4.21 | 0.03 |
| Peak frequency middle note | 0.16 | 0.4 |
| Peak frequency whole trill | 1.07 | 0.5 |
| Peak frequency lowest note | 0.77 | 0.49 |
| Peak frequency highest note | -1.37 | 0.49 |
| Peak frequency note 1 | 0.53 | 0.48 |
| Peak frequency note 2 | 1.35 | 0.49 |
| Peak frequency note 3 | 1.28 | 0.49 |
| Peak frequency note 4 | -1.32 | 0.39 |
| Peak frequency note 5 | -1.01 | 0.34 |
| Peak frequency 4th last note | -0.24 | 0.5 |
| Peak frequency 3rd last note | 0.67 | 0.5 |
| Peak frequency 2nd last note | -0.16 | 0.48 |
| Peak frequency last note | -0.93 | 0.43 |

**Table S3:** GLMMs with Gaussian distribution run in lme4 in R on experiment subset from May 2012, testing for differences between subject and playback stimulus singer in a) peak frequency (frequency differential) and b) body mass (body mass differential) that males and females do and do not respond to. Territory identity was a random factor in the models. (Please note the discrepancy between the samples sizes of female responses in the frequency and body mass tests is because one female was never recorded singing, so we had no value for her frequency to calculate differences with).

| **a) frequency differentials between experiments with and without a response** | **Estimate** | **SE** | **Z** | **P** |
|---|---|---|---|---|
| 1) Male songs females do and do not sing in response to (n = 16, territories = 8) | | | | |
| Intercept | -0.0001 | 0.007 | -0.02 | 0.99 |
| Female response | -0.03 | 0.01 | -3.02 | **0.01** |
| 2) Female songs females do and do not sing in response to (n = 16, territories = 8) | | | | |
| Intercept | -0.01 | 0.007 | -1.54 | 0.16 |
| Female response | 0.01 | 0.009 | 1.58 | 0.14 |
| 3) Male songs males do and do not sing in response to (n = 18, territories = 9) | | | | |
| Intercept | -0.01 | 0.006 | 1.89 | 0.08 |
| Male response | -0.01 | 0.007 | -1.98 | 0.06 |

| | | | | |
|---|---|---|---|---|
| 4) Females songs males do and do not sing in response to (n = 18, territories = 9) | | | | |
| Intercept | -0.003 | 0.01 | -0.32 | 0.75 |
| Male response | 0.006 | 0.01 | 0.51 | 0.62 |
| **b) body mass differentials between experiments with and without a response** | | | | |
| (all tests: n = 18, territories = 9) | **Estimate** | **SE** | **Z** | **P** |
| 5) Males that females do and do not sing in response to | | | | |
| Intercept | 0.001 | 0.01 | 0.14 | 0.89 |
| Female response | 0.04 | 0.02 | 2.65 | **0.02** |
| 6) Females that females do and do not sing in response to | | | | |
| Intercept | -0.001 | 0.01 | -0.12 | 0.91 |
| Female response | -0.02 | 0.02 | -0.80 | 0.44 |
| 7) Males that males do and do not sing in response to | | | | |
| Intercept | -0.01 | 0.01 | -0.39 | 0.70 |
| Male response | 0.02 | 0.01 | 1.19 | 0.26 |
| 8) Females that males do and do not sing in response to | | | | |
| Intercept | 0.02 | 0.02 | 0.72 | 0.48 |
| Male response | -0.03 | 0.02 | -1.28 | 0.23 |
